# Detecting Mitochondrial Free Radicals with Quantum Sensors: From Organelles to a *D. melanogaster* Model of Neurodegeneration

**DOI:** 10.1101/2025.11.29.691160

**Authors:** Shalini Menon, Jacob Reed, Brad Ebanks, Muralidharan Shanmugham, Adam Brookfield, David Collison, Maabur Sow, Victoria James, Nigel P. Mongan, Nicoleta Moisoi, Fedor Jelezko, Lisa Chakrabarti, Melissa L Mather

**Affiliations:** Diamond Quantum Sensing Hub, Institute of Advanced Manufacturing, Faculty of Engineering, University of Nottingham, Nottingham, NG7 2RD, UK; Optics and Photonics Research Group, Faculty of Engineering, University of Nottingham, Nottingham, NG7 2RD, UK; School of Veterinary Medicine and Sciences, University of Nottingham, Sutton Bonnington, LE12 5RD, UK; EPSRC National Research Facility for EPR and Spectroscopy, Department of Chemistry, The University of Manchester, Manchester, M13 9PL, UK; Institute for Quantum Optics, Ulm University, Albert-Einstein-Allee 11 D-89081 Ulm, Germany; School of Pharmacy, De Montfort University, The Gateway, Leicester, LE1 9BH, UK

**Keywords:** Quantum sensing, Nitrogen-Vacancy Centres, Mitochondrial free radicals, Neurodegeneration, optically detected magnetic resonance

## Abstract

This study identifies a distinct free radical signature in a D. melanogaster model of Parkinson’s Disease, successfully differentiating Pink1-deficient flies from wild-type controls. This insight was achieved using a robust quantum sensing methodology for the selective detection of free radicals in biological systems. Our approach utilizes optically detected magnetic resonance (ODMR) and magnetic modulation (MM) protocols with nanodiamond nitrogen-vacancy (NV) centres. Selective identification is achieved using the spin probe TEMPOL, a cell-permeable superoxide dismutase 2 (SOD2) mimic that initially quenches the photoluminescence signal.

Upon scavenging free radicals like hydroxyl and superoxide, TEMPOL is converted to a diamagnetic adduct, restoring the contrast and thus enabling quantitative detection. The approach was first validated in a chemical system with radical generation confirmed by electron paramagnetic resonance (EPR) spectroscopy. It was then demonstrated across biological scales, from isolated mitochondria and whole glioblastoma cells to the Drosophila model. In these studies, high-resolution respirometry revealed distinct free radical signatures, and findings were compared with NV T1 relaxometry. This work provides new biophysical insight into mitochondrial dysfunction, demonstrating a distinct free radical signature in a neurodegeneration model and connecting it to specific metabolic states.

## Introduction

Mitochondria, essential organelles in almost all eukaryotic cells, generate energy through oxidative phosphorylation (OXPHOS), a process intrinsically linked to the production of free radicals. These highly reactive and dynamic molecules are crucial for cellular signalling but can lead to oxidative damage when their production is dysregulated, associating with ageing, obesity, and various pathologies including neurodegeneration, cancer, and metabolic syndrome ^1^. Understanding the precise relationship between mitochondrial function and free radical dynamics is, therefore, critical for both fundamental biological insights and clinical advancements.

Current methodologies for assessing mitochondrial function and free radical production often rely on measuring stable by-products of oxidative stress or employing electron paramagnetic resonance (EPR) spectroscopy with spin probes or traps. While EPR offers specificity for paramagnetic species, its sensitivity is often limited in biological contexts, particularly at low radical concentrations ^2^. Recent advancements in EPR technology have enhanced sensitivity but frequently introduce increased complexity and reduced accessibility ^3^.

Quantum sensing, particularly utilizing nitrogen-vacancy (NV) centres in diamond, presents a promising alternative for highly sensitive detection of paramagnetic species such as free radicals. Among the diverse NV centre-based sensing protocols, optically detected magnetic resonance (ODMR) is widely employed. Continuous wave (CW) ODMR uses simultaneous continuous green light illumination and microwave frequency sweeps to detect changes in NV photoluminescence (PL) intensity at resonant frequencies. Quantum decoherence protocols (sensing approaches employing change of coherence time as a signal), employing pulsed green light and microwaves, are used to measure changes in longitudinal (T_1_) and transverse (T_2_) relaxation times to detect target analytes. T_1_ relaxometry is notably prevalent for detecting paramagnetic species like free radicals, as their presence shortens NV T_1_ times. Significant advancements include the sensitive characterization of radical species and spatio-temporal mapping of redox reactions in aqueous solutions ^4^, and the detection of free radicals in individual yeast cells under various conditions ^5^. Schirhagl and colleagues have pioneered the use of T_1_ relaxometry for intracellular free radical detection, highlighting its potential in biological applications 6–13.

However, while T1 relaxometry provides quantitative information, it demands precise hardware synchronization and a light source that can rapidly switch, has excellent power stability and free from high-frequency noise. Moreover, photoionization of the NV charge state during measurement can act as an experimental artefact, further complicating data interpretation^11,12^ . Inherent variability in T_1_ times when using diamond particles can also lead to measurement uncertainties, requiring longer acquisition times and making it less suitable for tracking highly transient species. Alternative approaches have leveraged PL differences between NV charge states to quantify hydroxyl radicals ^13^ or used coupled charge and spin dynamics for sensitive detection ^14^. Despite these innovations, challenges persist in terms of technical complexity and accessibility.

The present study reports an innovative and accessible quantum sensing approach for selective free radical detection in biological systems. Our methods harness the spin-dependent photoluminescence (PL) of negatively charged NV centres in nanoscale diamonds, employing simple modifications to a standard inverted fluorescence microscope operating at low optical power to preserve the NV charge state while maintaining sensitivity to paramagnetic species. We utilize continuous-wave optically detected magnetic resonance (CW ODMR) and a microwave-free magnetic modulation (MM) method, which leverages the dependence of NV PL on an applied magnetic field. Building on previous work, this study demonstrates that the extent of PL contrast quenching induced by a magnetic field can be used for the detection of paramagnetic species, particularly free radicals.

Furthermore, this study employs the spin probe TEMPOL (4-hydroxy-2,2,6,6-tetramethylpiperidine-N-oxyl), a cell-permeable mimic of superoxide dismutase 2 (SOD2), for selective free radical identification. Initially, TEMPOL serves as a control by quenching both ODMR and MM photoluminescence contrast. The subsequent restoration of this contrast upon TEMPOL’s reaction with radicals like hydroxyl and superoxide provides a quantitative measure of their production. This methodology was first validated in a chemical system (UV photolysis of H₂O₂), with free radical generation confirmed by EPR spectroscopy. For benchmark data and cross-comparison, our NV measurements were performed alongside EPR, NV T_1_ relaxometry, and high-resolution respirometry (Oroboros O2K) protocols on isolated mitochondria. The approach was then applied to a spectrum of biological systems, ranging from isolated mitochondria and whole glioblastoma cells to *D. melanogaster*. We focused on the *Pink1*-deficient *D. melanogaster*, a key model for early-onset Parkinson’s Disease (PD) known to accumulate damaged mitochondria and elevated Reactive Oxygen Species (ROS) ^15,16^.

By integrating our sensing protocols with specific substrate-uncoupler-inhibitor titrations (SUITs) in high-resolution respirometry, we demonstrate the ability to distinguish between control (wild-type) and diseased (*Pink1*-deficient) organisms. This showcases a powerful new tool for investigating the biophysical parameters underlying mitochondrial dysfunction in disease. This accessible technique significantly lowers the barrier to entry for quantum sensing applications in biology, providing a unique window into the complex interplay of energy and free radical production crucial for health and disease, with immediate applications in cellular toxicity evaluation, drug screening, and fundamental research.

## Materials and Methods

### Cell Culture

The paediatric glioblastoma cell line SF188 (Sigma Aldrich) was cultured in T75 or T175 flasks (for whole-cell analysis or mitochondrial isolation, respectively) using Dulbecco’s Modified Eagle Medium (DMEM) (Sigma Aldrich) supplemented with 10% Fetal Bovine Serum (FBS) (Sigma Aldrich) and L-glutamine (Sigma Aldrich). Cells were maintained at 37°C with 5% CO_2_ and harvested at 90% confluency.

### Isolation of Mitochondria (Mt)

Enriched mitochondrial fractions were isolated by differential centrifugation as previously described^17^. Briefly, cell pellets were resuspended in mitochondrial extraction buffer (50 mM Tris-HCl pH 7.4, 100 mM KCl, 1.5 mM MgCl_2_, 1 mM EGTA, 50 mM HEPES, 100 mM sucrose; Sigma-Aldrich) and homogenized using a GentleMACS Dissociator (Miltenyi Biotec).

Homogenates were centrifuged sequentially at 4°C: 850 × g (10 min), 1000 × g (10 min, to pellet nuclei), and 10,000 × g (10 min) to obtain the mitochondrial pellet.

### D. melanogaster Husbandry

Fly stocks and crosses were maintained in Blades Biological *D. melanogaster* Quick Mix Medium (Blue) at 24° C in a 12:12 light-dark cycle. Experiments were performed on male flies: wildtype referred to as WT (genotype *w1118*) and Pink1^-^ (genotype *Pink1B9/Y*). The flies were transferred to new food every 3 days and were prepared by homogenization, for respirometry in the Oxygraph-2k (Oroboros Instruments, Innsbruck, Austria) at 25 days post-eclosion.

### High-Resolution Respirometry (HRR)

Mitochondrial physiology was assessed using an Oxygraph-2k FluoRespirometer (Oroboros Instruments, Innsbruck, Austria). High resolution respiratory data was collected at 37°C. All experiments were performed in mitochondrial respiration medium (MiR05, Oroboros Instruments - consisting of 0.5 mM EGTA, 3 mM MgCl₂, 60 mM lactobionic acid, 20 mM taurine, 10 mM KH₂PO₄, 20 mM HEPES, 110 mM D-sucrose, and 1 g/L BSA, pH 7.1), with a constant stirring speed of 750 rpm. An air calibration was performed prior to each experiment (www.bioblast.at).

To minimize interference with cellular respiration while ensuring sufficient quenching of NV photoluminescence contrast, increasing concentrations of TEMPOL were tested on whole-cell oxygen consumption rates (OCRs). These concentrations were evaluated in, parallel temperature controlled tubes outside the respirometer chambers to determine the optimal level. A concentration of 25 mM TEMPOL was selected as it did not alter OCRs and provided reversible quenching within a biologically relevant free radical concentration range.

For experiments with isolated mitochondria and then with whole cells, samples were suspended in 2 mL of MiR05 buffer containing 25 mM TEMPOL and added to the respirometer chamber. A substrate-uncoupler-inhibitor (SUIT) protocol, depicted in Supplementary Figure S1, (SUIT-004 O2 pfi D010) was then initiated with the sequential addition of pyruvate (final concentration 5 mM) and malate (2 mM), followed by titrations of carbonyl cyanide m-chlorophenyl hydrazone (CCCP; 0.5 µM). Finally, rotenone (2.5 µM) and antimycin A (2.5 µM) were added to inhibit the electron transport chain proteins Complex I and Complex III, respectively.

For *D. melanogaster* studies, a different SUIT protocol was used. Three whole flies were homogenized in 300 µL of MiR05, and 200 µL of the supernatant was added to each chamber. After permeabilization with digitonin (10 µg/mL), the following substrates and inhibitors were added sequentially: pyruvate (5 mM) and proline (10 mM), ADP (2 mM), cytochrome C (10 µM), succinate (10 mM), titrations of CCCP (0.125 µM), rotenone (0.5 µM), and antimycin A (2.5 µM). Oxygen consumption and flux were recorded continuously throughout both protocols.

### Nanodiamond Preparation

Carboxylated red fluorescent NDs (FNDBiotech, Taiwan), 100 nm nominal diameter with >1000 NV centres per particle, were used for all NV sensing experiments. Glass coverslips (No. 1.5, 18 mm x 18 mm) were coated with NDs via drop-casting of 2.5 μL of a 0.2 mg/mL ND suspension in deionized water, followed by overnight drying at 60°C in an oven. The nanodiamond-coated coverslips were stored in the dark at room temperature until use.

### Optical and Microwave Setup

The FND-coated coverslips were mounted onto a custom-designed printed circuit board (PCB), engineered for microwave (MW) delivery, and secured on the sample stage of an inverted fluorescence microscope (Olympus IX83), depicted in Figure 1. NV centres were excited using the 550 nm line of a CoolLED pE-4000 Illumination System, filtered through a 545 ± 15 nm bandpass filter, delivering approximately 0.5 mW to the sample plane over a spot size of approximately 220 µm in diameter. Light was focused onto the sample using an Olympus UPlanxAPO 60× oil-immersion TIRF objective lens (NA = 1.49). The resulting PL was collected by the same objective, passed through a longpass filter (575 nm cut on wavelength) and imaged onto a sCMOS camera (Photometrics Prime 95B). All hardware control and data acquisition were managed using Olympus CellSens software version 2.0 and an Olympus Real-Time Controller (RTC).

**Figure 1.**
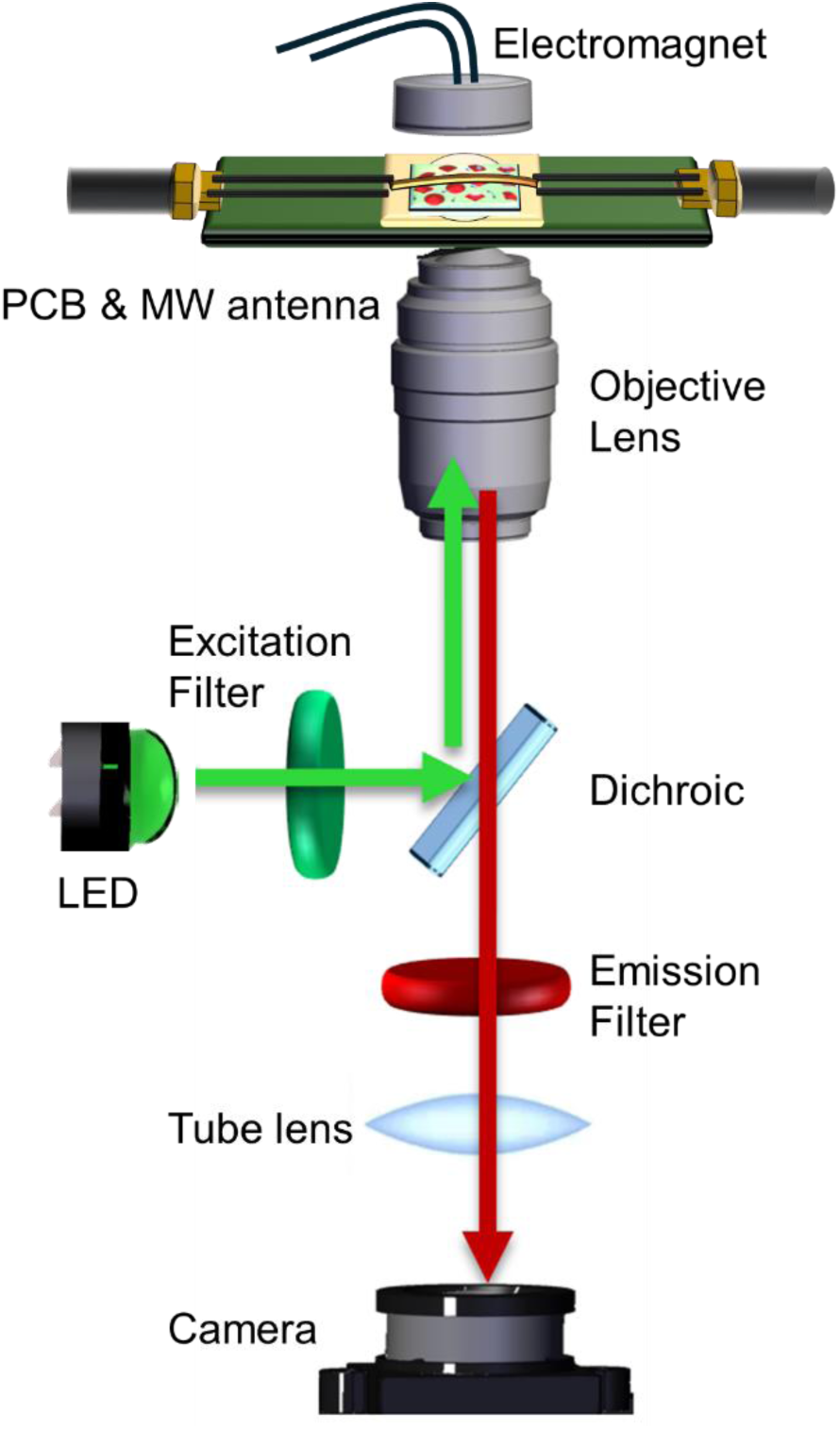
A schematic representation of the NV sensing instrumentation is shown. Microwave delivery is achieved via a PCB-coupled wire, while an electromagnet provides amplitude-modulated magnetic fields. Optical detection is performed using an inverted fluorescence microscope equipped with green LED illumination, excitation filter, dichroic, an oil-coupled objective for light delivery and collection, an emission filter, tube lens and an sCMOS camera for image acquisition. A pipette is used for delivery of the sample to a nanodiamond coated microscope glass coverslip fixed to the PCB.

### Continuous-Wave ODMR

Continuous-wave (CW) ODMR spectra were acquired using an Agilent E4428C signal generator (0 dBm output), amplified by an AR 20S1G4 MW amplifier (30 % gain). Microwaves were delivered (input power of 1 W) to the FNDs via a 0.125 mm diameter copper wire on the PCB. For each spectrum, the microwave frequency was swept from 2.77 GHz to 2.97 GHz in 2 MHz steps, with a camera exposure of 30 ms per step, generating an image stack where each frame corresponds to a specific microwave frequency. Each full sweep was repeated 10 times for signal averaging.

### Magnetic Modulation (MM)

For MM measurements, an external magnetic field of 40 ± 1 mT was applied using an electromagnet (verified by a Hall probe) driven by a Tektronix AFG 3102 function generator. The field was square-wave modulated at 0.5 Hz (a 2-second cycle with 1 s on, 1 s off). Data was acquired continuously over 4 modulation cycles (8.0 s total duration). A total of 210 frames were captured in each experiment, with each frame having an exposure time of 30 ms and an additional readout/processing time (∼ 8 ms), resulting in an effective frame rate of approximately 38.1 ms per frame. Data were acquired continuously over 4 modulation cycles, capturing a total of 210 frames with a 30 ms exposure time per frame, saved as an image stack. This protocol allowed for robust averaging of the PL signal in both the “field on” and “field off” states.

### Image Processing and Region of Interest (ROI) Selection

All image stacks from ODMR and MM acquisitions were processed using ImageJ software (v. 1.54). From a representative image in each stack, multiple regions of interest (ROIs), approximately 10 µm in diameter and corresponding to isolated FND clusters, were manually selected. For each ROI, the average PL was calculated by averaging the pixel intensities within its boundary. This average PL value for each individual ROI was then tracked across the entire image stack. This process yielded distinct datasets depending on the measurement type: for ODMR, a plot of PL as a function of the applied microwave frequency, and for MM, a plot of PL corresponding to the ’magnetic field on’ or ’magnetic field off’ state of each frame.

### ODMR and MM Contrast Calculation

For ODMR analysis, the PL intensity dataset for each ROI was processed using OriginLab software (version 2022). The data were first normalized to the average intensity of the initial off-resonance frames (2.77 GHz –2.80 GHz). A double Lorentzian function was then fitted to the normalized spectrum to precisely determine the on-resonance and off-resonance PL values. The ODMR contrast was defined as the normalized difference between these two fitted values. For MM analysis, the contrast was calculated as the normalized difference between the average PL intensity measured during the ’field on’ state and the ’field off’ state across the full acquisition.

### Normalization and Statistical Analysis

To enable comparison across different biological samples and experiments, a two-step normalization procedure was applied to the calculated contrast values. First, the contrast from a given sample was normalized to the baseline contrast measured from a 25 mM TEMPOL control solution. Second, to specifically track the restoration of the signal due to free radical scavenging, the contrast change within an experiment was normalized relative to that sample’s own starting value (i.e., post-TEMPOL addition, pre-stimulus). In this final representation, a value of 1.0 signifies maximum signal quenching by TEMPOL, with values >1.0 indicating contrast restoration due to radical production. The number of replicates for each experiment is detailed in the relevant figure captions. Data are presented as mean ± standard error of the mean apart from T_1_ relaxation times which are presented as standard deviation obtained from fitting the T_1_ measurement data to an exponential decay function.

### Free Radical Calibration via H₂O₂ Photolysis

A calibration curve correlating hydroxyl radical (OH•) concentration with normalized ODMR and MM contrast was established. Stock solutions of 10%, 20%, and 30% (v/v) hydrogen peroxide (H_2_O_2_) (Sigma-Aldrich, UK) were prepared in Milli-Q water. For each calibration point, an aqueous mixture containing 25 mM TEMPOL (Sigma-Aldrich, UK) and one of the H₂O₂ concentrations was prepared. OH• radicals were generated by irradiating these mixtures for 3 minutes with 365 nm UV light from a CoolLED pE-4000 Illumination System at 100% power setting. Immediately following irradiation, a 100 µL aliquot of the mixture was pipetted onto an FND-coated coverslip, and ODMR and MM data were recorded. The concentration of OH• radicals produced under these conditions was independently quantified in parallel experiments using EPR spectroscopy with a spin trap (see EPR Validation Studies section and Supplementary Figure S2)

### EPR Validation Studies

EPR spectroscopy was employed to characterize radical species generated under various conditions. Two sets of samples were prepared. First, radical species from the UV photolysis of aqueous solutions containing 25 mM TEMPOL and H₂O₂ were analysed. Second, to examine radical production during cellular respiration, mitochondrial and whole-cell samples were incubated with either 2.5 mM TEMPOL or 50 mM of the spin trap 5,5-dimethyl-1-pyrroline N-oxide (DMPO). The lower TEMPOL concentration (2.5 mM) was selected to allow for monitoring changes in the intensity of its characteristic three-line nitroxide signal, while the 50 mM DMPO concentration was used to identify the specific radical species generated. Following preparation, all samples were flash-frozen in liquid nitrogen and stored at −80°C until analysis.

EPR analyses were conducted following established protocols. Immediately before measurement, thawed samples were transferred into 1.3 mm outer diameter (OD) / 1 mm inner diameter (ID) glass capillary to a sample height of approximately 40 mm. Each capillary was inserted into a 4 mm OD / 3 mm ID quartz EPR tube (Wilmad LabGlass) and positioned within a Bruker ER 4112SHQ X-band resonator for optimal signal intensity.

All EPR spectra were acquired at room temperature on a Bruker EMXmicro EPR spectrometer with the following parameters: microwave power of 1 mW (23 dB), modulation amplitude of 0.5 G, time constant of 82 ms, conversion time of 1 ms, sweep time of 30 s, receiver gain of 30 dB, and an average microwave frequency of 9.851 GHz. The resulting continuous wave EPR spectra were analysed and simulated using the EasySpin (v. 5.2.35) toolbox for MATLAB toolbox^18^.

### T1 Relaxometry Comparison Measurements

T_1_ relaxometry measurements were performed on isolated mitochondria samples at each of the six stages of the SUIT protocol. Measurements were performed with a custom made confocal set up described in a previous publication^19^. To enhance measurement fidelity, a pre-screening step was first conducted on nanodiamonds in air to identify diffraction limited FND clusters (less than 500 nm) with the longest intrinsic T_1_ times. A long intrinsic T_1_ provides a broader dynamic range for detecting T_1_ shortening; this ensures that even after the relaxation is accelerated by radicals, the decay curve can still be sampled robustly across multiple time points, improving the precision and reliability of the final fitted T_1_ value. A single, pre-characterized FND was then used for the entire measurement sequence. Before each new solution was added, the sample well was rinsed three times to prevent cross-contamination.

For NV spin initialization and readout, a green laser (513 nm Toptica Ibeam smart 515.S-15133) was used, resulting in a power of 20 µW at the sample plane. An alternating T_1_ pulse sequence was employed as previously reported with the exception that microwave pulses were rapid adiabatic pulses instead of square one^20,21^. The robust coherent control offered by rapid adiabatic pulses improves measurement contrast and do not require bias magnetic field^22^. The delay time (τ) between laser pulses was varied logarithmically over 10 points from 1 µs to 3 ms. The microwave pulses, amplified by the 16W amplifier (250 mV peak amplitude in the AWG), had a duration of 10 µs a frequency range of 2.83 GHz to 2.91 GHz and a shape of the Allen-Eberly model with a truncation ratio of 0.1^23^. Each T_1_ measurement was acquired rapidly (approximately 3 minutes). The resulting photoluminescence decay curves were fitted to a stretched exponential function to extract the T_1_ value.

## Results and Discussion

### Demonstration of Free Radical Detection using H₂O₂ Photolysis

The functionality of the TEMPOL-based quantum sensing approach was first validated using UV photolysis of H₂O₂ to generate hydroxyl radicals (OH•). As shown in figures 2a and 2b, the addition of 25 mM TEMPOL alone significantly reduced the baseline ODMR and MM contrast compared to nanodiamonds in air, which is consistent with paramagnetic quenching from the TEMPOL probe. Upon UV irradiation of H₂O₂ solutions containing TEMPOL, this contrast was restored. This observation supports the hypothesis that TEMPOL scavenges the generated OH• radicals and is converted to a diamagnetic, non-quenching species, demonstrating the viability of the sensing mechanism. Figure 2c shows that the extent of this contrast restoration is quantitatively dependent on the initial H₂O₂ concentration. Increasing the H₂O₂ concentration from 0% to 30% resulted in a recovery of the ODMR contrast from approximately 2.5% to 5.5%, and the MM contrast from approximately 6% to 10.8%. As shown in Supplementary Figure S2, varying concentrations of DMPO-OH and DMPO-R adducts are produced when H_2_O_2_ was irradiated with UV-LED (365 nm) at room temperature. This allowed the creation of a calibration curve that correlates the NV sensor’s contrast with radical concentration, establishing that the ODMR and MM contrast can be used to measure relative changes in free radical concentration. While prior work has focused on real-time detection via changes in the NV charge state^9,24^, our method enables offline, selective detection of radical concentrations using the TEMPOL stabilising probe in sample aliquots^25^.

**Figure 2:**
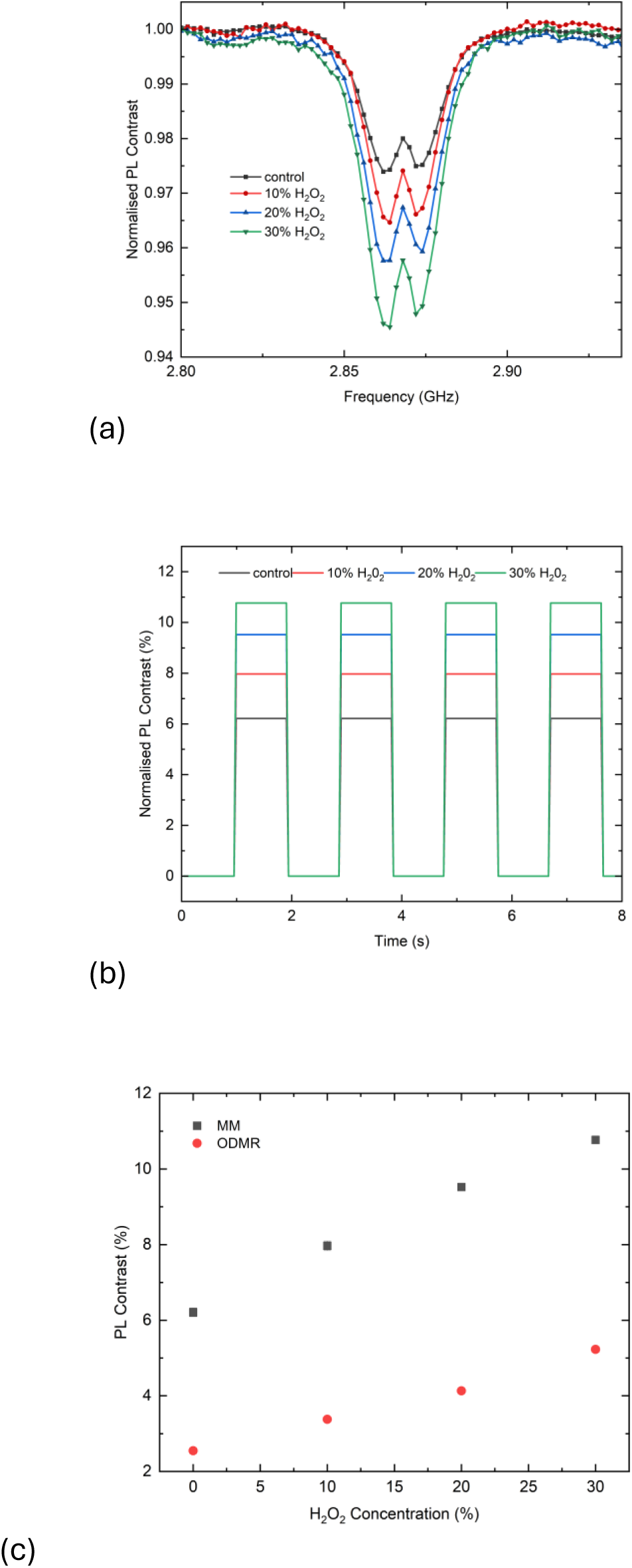
TEMPOL-mediated Modulation of ODMR and MM Contrast in the Presence of Hydroxyl Radicals. (a) ODMR spectra demonstrating the effect of 25 mM TEMPOL on NV centres in nanodiamonds (NDs) following UV photolysis of varying H₂O₂ concentrations (irradiation time: 3 minutes). (b) Corresponding MM spectra under the same conditions as (a). The crest represents the average of “OFF” values and the trough the average of “ON” values across the frames. (c) Comparison of PL ODMR and MM contrasts obtained from various H₂O₂ concentrations with the 25 mM TEMPOL control. (n = 5 replicates)

### Free Radical detection Across Biological Scales: From Isolated Mitochondria to Whole Organisms

The diamond sensor was first applied to detect free radical production in isolated mitochondria and whole SF188 glioblastoma cells during a SUIT protocol (SUIT004D010). For both sample types, a progressive increase in ODMR and MM contrast was observed throughout the SUIT protocol (Figure 3). These trends are summarized by the contrast values plotted for ODMR and MM data in Figures 3a and 3b, respectively. In isolated mitochondria, the ODMR contrast increased from an initial ∼6.7% to a maximum of ∼7.5%, while the MM contrast increased from ∼6.9% to ∼7.4%. In whole cells, the ODMR contrast increased from ∼5% to ∼6%, and MM contrast from ∼9.6% to ∼10.4%. These rising contrast values indicate the ongoing consumption of the paramagnetic TEMPOL probe, consistent with its reaction with mitochondrially-generated free radicals. These trends are also illustrated in the underlying ODMR spectra (Figures 3c and e) and MM traces for mitochondria and cells (Figures 3d and f), respectively.

**Figure 3:**
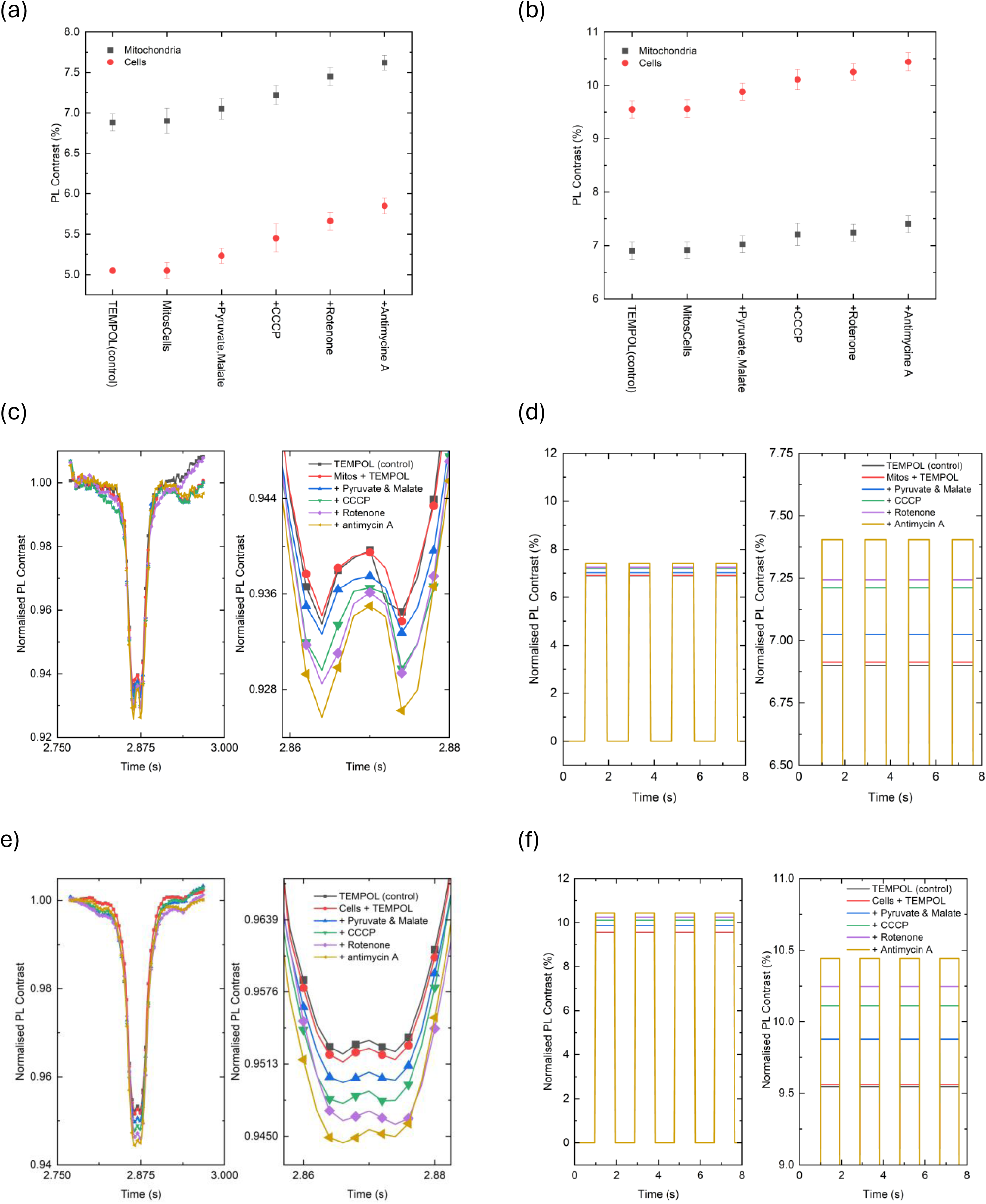
Photoluminescence (PL) contrast derived from ODMR and MM measurements is shown for isolated mitochondria and whole cells (a, b), No. of replicates = 5. Normalized ODMR contrast and MM contrast are presented for mitochondria throughout the SUIT protocol (c, d), and for whole cells throughout the SUIT protocol (e, f).

The observed changes in sensor contrast corresponded to specific metabolic states induced by the SUIT protocol. In both sample types, contrast increased upon the addition of substrates (pyruvate and malate), a finding that aligns with reported OH• production, and further increases with the uncoupler CCCP are consistent with enhanced radical production due to uncoupled respiration. Subsequent increases in contrast following the inhibition of Complex I (rotenone) and Complex III (antimycin A) align with the known phenomenon of elevated free radical production under these conditions (Figure 4). These trends were directly compared with the mitochondrial respiratory physiology by calculating the Flux Control Ratio (FCR) from the respirometry data, which normalizes the oxygen consumption rate at each metabolic state to the maximum respiratory capacity (Figure 4). This comparison showed that while the ODMR and MM contrast consistently increased across all states, indicating cumulative free radical production, the FCR exhibited a more dynamic trend, decreasing upon the inhibition of Complex I and Complex III. This finding suggests that the level of free radical production does not simply scale with the rate of mitochondrial oxygen consumption. Instead, a breakdown in mitochondrial function, as seen with respiratory chain inhibitors, leads to a decrease in overall respiratory rate but a continued increase in the generation of free radicals.

**Figure 4:**
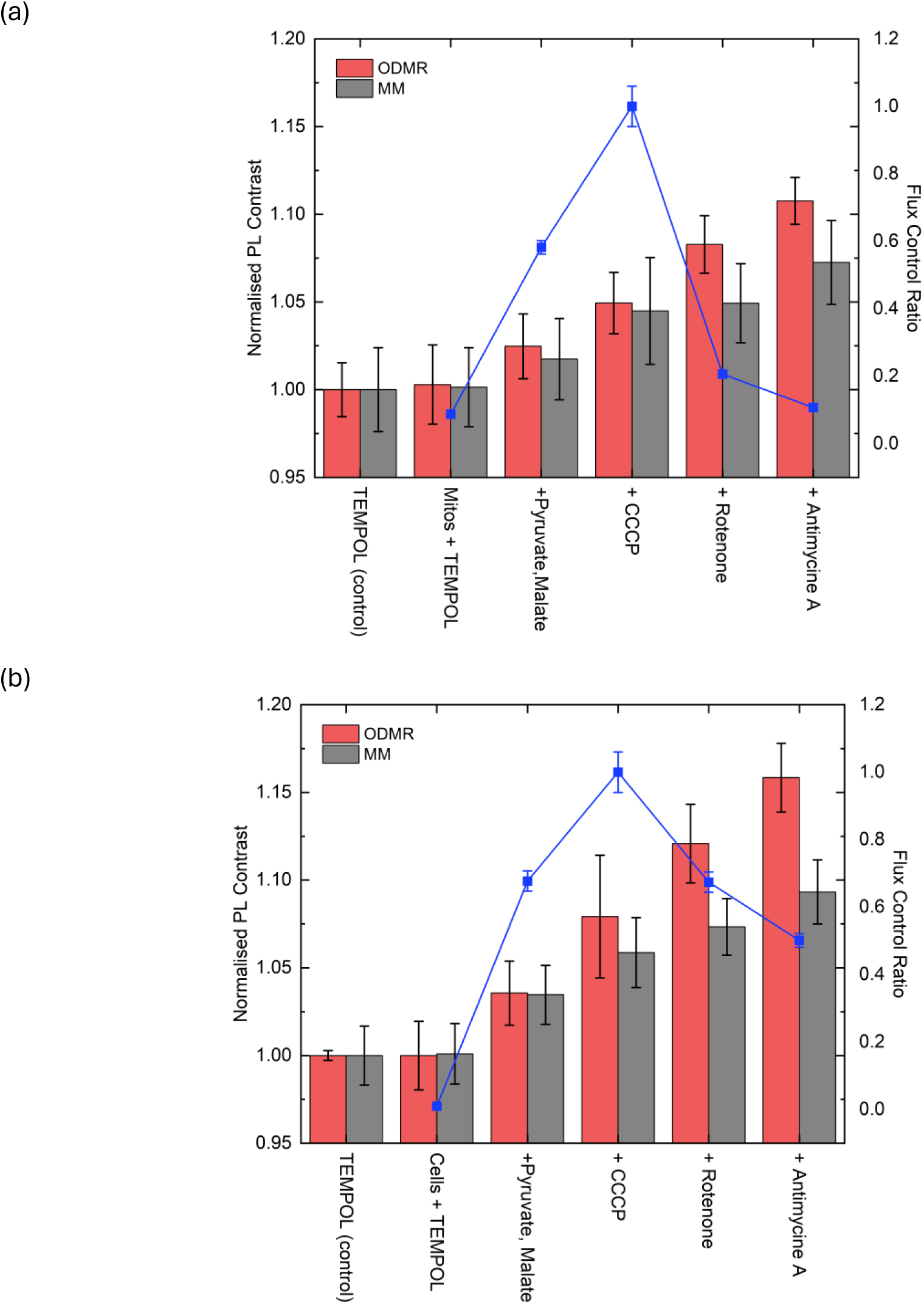
Comparison of the Flux Control Ratio (FCR, blue line) values (Right y-axis, blue points and line) with the normalised ODMR and MM contrasts derived from the contrast ratio of isolated mitochondria to 25 mM TEMPOL (control) and whole cells to 25 mM TEMPOL (control) No. of replicates = 5

These findings were corroborated by multiple validation techniques. T_1_ relaxometry measurements performed on the isolated mitochondria showed a corresponding recovery of the T_1_ time from ∼0.5 ms to ∼1.5 ms during the titration, Supplementary Figure 3. Furthermore, EPR spectroscopy on whole-cell samples—treated with TEMPOL, pyruvate, malate, and CCCP—provided direct evidence for the proposed mechanism. A significant (∼80%) decrease in the TEMPOL nitroxide signal was observed, supporting its consumption by reaction with free radicals (Figure 5a). Finally, spin-trapping experiments with DMPO confirmed the identity of the species being produced, revealing a characteristic 1:2:2:1 intensity pattern of the DMPO-OH adduct and confirming hydroxyl radical generation during cellular respiration (Figure 5b). The observed hyperfine coupling to the *a*_iso_(^14^N) = 39.7 MHz and *a*_iso_(^1^H) = 33.6 MHz (simulated datum not shown) nuclei for this adduct aligned with previously reported values^26^.

**Figure 5:**
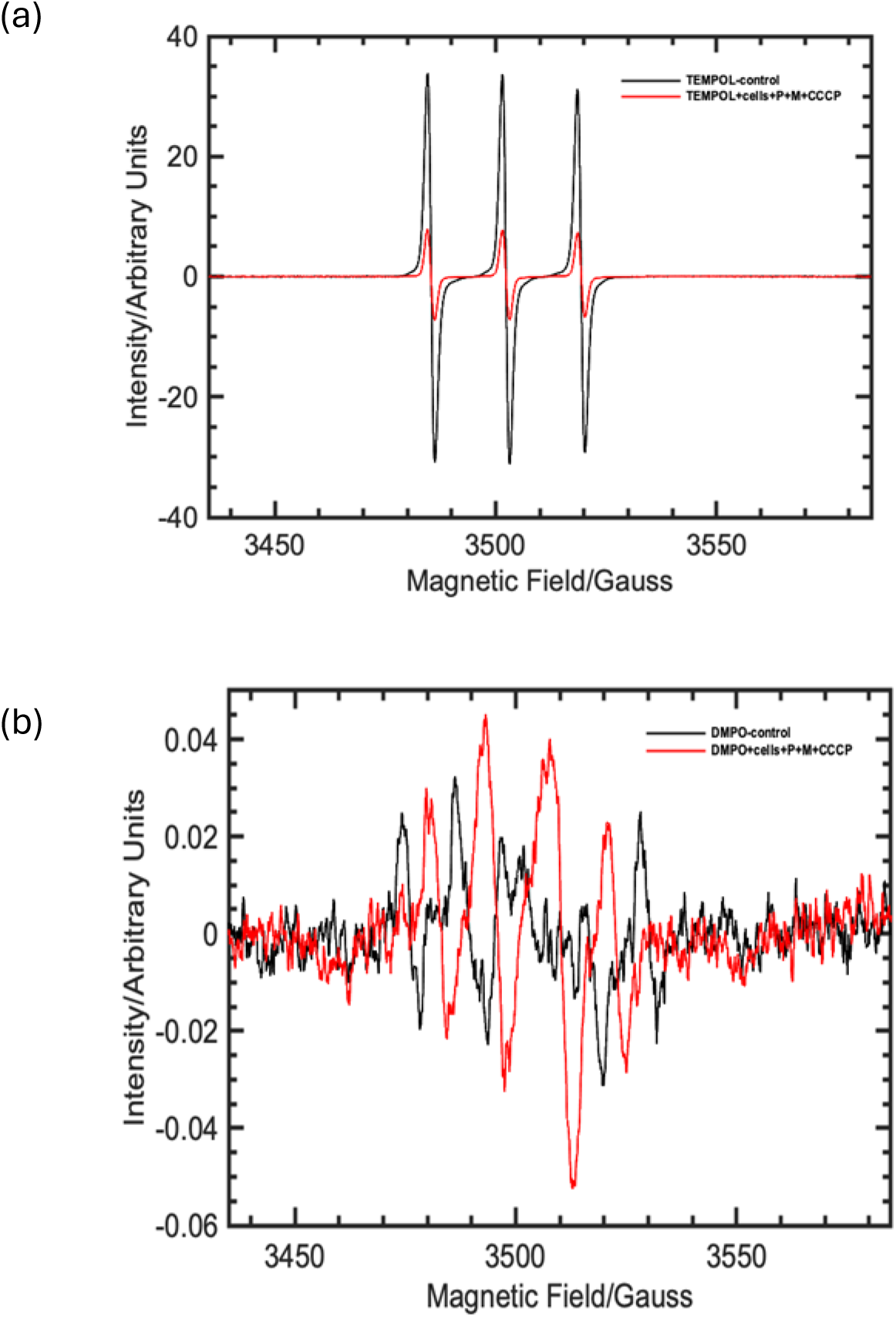
(a) Continuous wave EPR spectra of 2.5 mM TEMPOL in the absence (control; black trace) and presence of cells+substates+uncoupler (red trace), (b) Continuous wave X-band EPR spectra of 50 mM DMPO in the absence (control; black trace) and presence of cells+substates+uncoupler (red trace).

Building on these cellular and subcellular findings, the methodology was then applied to a whole organism model, comparing wild-type (WT) *D. melanogaster* to the *Pink1-deficient* mutant, a model for Parkinson’s Disease. Inactivating mutations affecting Pink1, the Pten-induced kinase-1 protein, regulates mitochondrial morphology, mitophagy and neuronal viability, thus contributing to Parkinson’s Disease^27^. This approach achieves chemical selectivity by utilizing the spin probe TEMPOL, which is known to preferentially scavenge reactive oxygen species such as OH• and superoxide radicals, enabling the quantitative tracking of relative changes in their concentration within the sensing volume. The primary finding, shown in Figure 6, was that *Pink1-deficient* flies exhibited a greater restoration of ODMR and MM contrast throughout the SUIT protocol compared to their wild-type counterparts, indicative of higher net free radical generation. Quantitatively, the ODMR contrast in *Pink1* flies increased from ∼4.7% to 5.6% (compared to ∼4.7% to ∼5.5% in WT), and the MM contrast increased from ∼9.7% to 11.3% (compared to ∼8.9% to ∼9.9% in WT). Analysis of the high resolution respirometry data suggests that the basal level of free radical formation is higher in the *Pink1-mutant* model and that a substantial increase in free radicals is observed in the uncoupled state, suggesting that *Pink1* deficiency may lead to mitochondrial dysfunction when the electron transport chain operates at maximum capacity.

**Figure 6:**
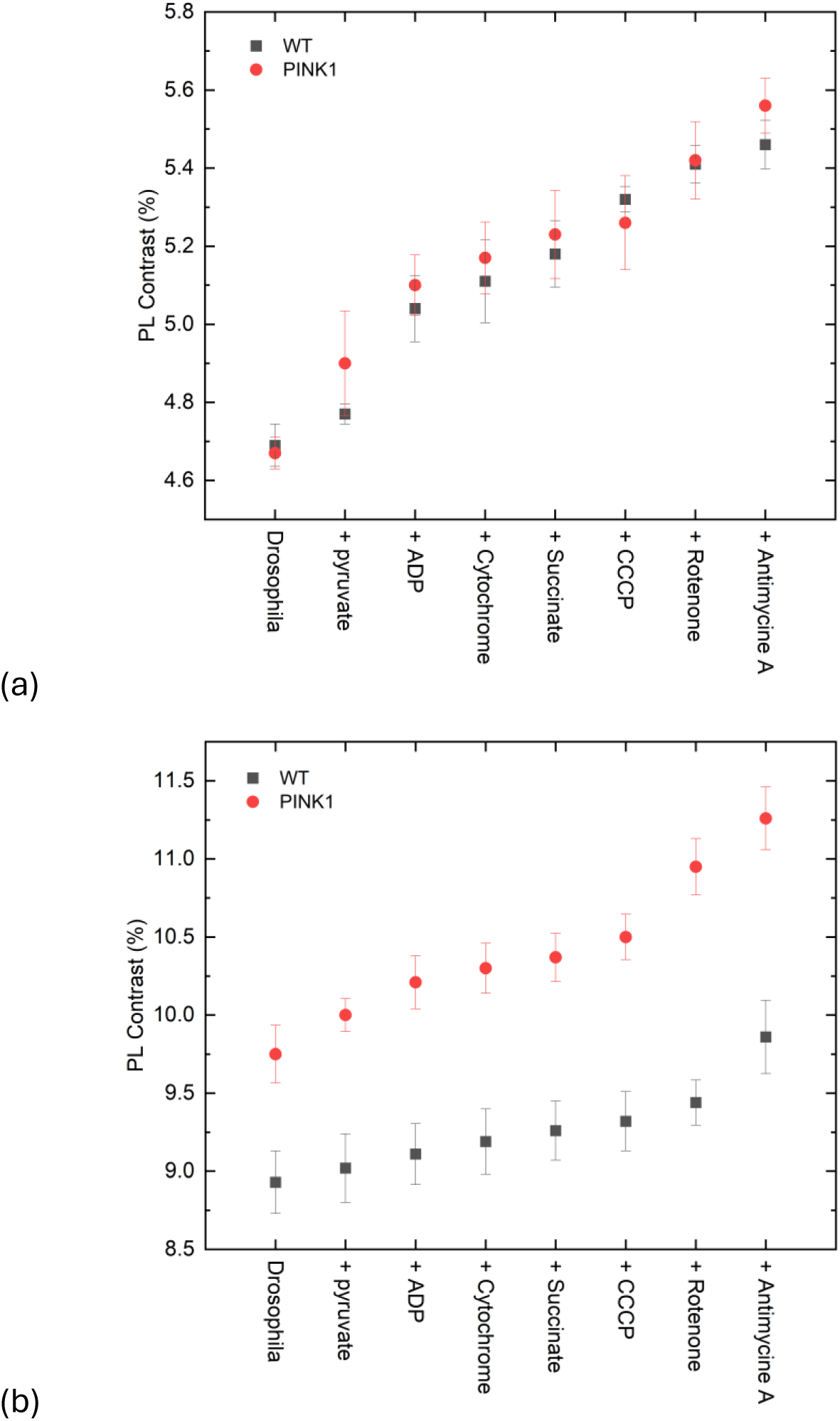
Photoluminescence (PL) contrast derived from ODMR (a) and MM (b) measurements is shown for wild-type (WT) *D. melanogaster* and the *Pink1*-deficient mutant model, No. of replicates = 5. *Pink1*-deficient flies exhibited a greater restoration of ODMR and MM contrast throughout the SUIT protocol compared to their wild-type counterparts, indicative of higher net free radical generation.

Collectively, these NV-based measurements align with observations from mitochondrial respiratory physiology. Notably, throughout the respiratory pathway, the whole SF188 cells exhibited larger changes in ODMR and MM contrast compared to the isolated mitochondria. The findings in the *D. melanogaster* model are consistent with the known biology of *Pink1*; it has been reported that ROS accumulation recruits *Pink1*/Parkin to initiate mitophagy, a process impaired in *Pink1-mutants* ^28^. The resulting ROS buildup in dysfunctional mitochondria may therefore explain the higher basal and stress-induced free radical levels detected in this work. This technique allows mitochondrial respiratory physiology to be compared directly to free radical formation at specific states of respiration in tandem, providing insights into homeostatic mechanisms of mitochondrial function at the quantum level.

## Conclusion

In this study, we have developed and validated a robust quantum sensing platform for the quantitative tracking of relative changes in free radical concentrations across biological scales. By combining nanodiamond NV-based measurements with the TEMPOL spin probe, we achieved sensitive detection of free radicals linked to mitochondrial respiration. The key significance of this work is demonstrated by its application to a *D. melanogaster* model of neurodegeneration, where the method was able to clearly distinguish between the elevated free radical production in *Pink1*-deficient mutants and their wild-type counterparts, particularly under conditions of maximal respiratory stress. This finding, differentiating between healthy and diseased states in a whole organism, represents a significant advance and highlights the platform’s potential for investigating the complex interplay between mitochondrial dysfunction and disease pathology.

The ability to align these sensing measurements with metabolic states mapped by high-resolution respirometry provides a powerful new tool for applications ranging from fundamental studies of oxidative stress to drug screening and toxicology. Looking forward, this work paves the way for more efficient analyses of radical-related disease processes. Future work will focus on employing spin probes specific to other radical types to further enhance the technique’s specificity and broaden its application in biomedical research.

## Author Contributions

S.M. performed the NV sensing experiments, data analysis, and EPR sample preparation. S.M., J.R., and B.E. conducted the biological work and respirometry. M.S. (Shanmugham) performed and analysed the EPR experiments under the supervision of A.B. and D.C. M.S. (Sow) performed and analysed the T1 experiments under the supervision of F.J. L.C. and M.L.M. conceived and designed the study. L.C. designed and oversaw the biological studies. M.L.M. designed the quantum sensing protocol and supervised the overall project. N.M. provided the Pink1 fly models. V.J. and N.P.M. assisted in securing funding. M.L.M. wrote the original manuscript draft. All authors reviewed and edited the manuscript.

## Data Availability

The datasets generated during and/or analysed during the current study are available from the corresponding author on reasonable request.

## Acknowledgements

This work was supported by funding from the Biotechnology and Biological Sciences Research Council (BBSRC) (grant number BB/T012226/1), the Royal Academy of Engineering through their Chair in Emerging Technologies Scheme (grant number CiET-2223-102), and the European Research Council (ERC) through the ERC Consolidator Award, TransPhorm (grant agreement ID: 683108). We acknowledge the EPSRC for access to the National Research Facility for EPR Spectroscopy (grant numbers EP/W012226/1, EP/S033181/1, EP/V035231/1, EP/X034623/1).

## References

1. Xu, X., Pang, Y. & Fan, X. Mitochondria in oxidative stress, inflammation and aging: from mechanisms to therapeutic advances. Signal Transduct Target Ther 10, 1–29 (2025).

2. Abbas, K., Babić, N. & Peyrot, F. Use of spin traps to detect superoxide production in living cells by electron paramagnetic resonance (EPR) spectroscopy. Methods 109, 31–43 (2016).

3. Ranjan, V. et al. Electron spin resonance spectroscopy with femtoliter detection volume. Appl Phys Lett 116, 184002 (2020).

4. Barton, J. et al. Nanoscale dynamic readout of a chemical redox process using radicals coupled with nitrogen-vacancy centers in nanodiamonds. ACS Nano 14, 12938–12950 (2020).

5. Morita, A. et al. Detecting the metabolism of individual yeast mutant strain cells when aged, stressed or treated with antioxidants with diamond magnetometry. Nano Today 48, 101704 (2023).

6. Wu, K. et al. Diamond Relaxometry as a Tool to Investigate the Free Radical Dialogue between Macrophages and Bacteria. ACS Nano 17, 1100–1111 (2023).

7. Perona Martínez, F., Nusantara, A. C., Chipaux, M., Padamati, S. K. & Schirhagl, R. Nanodiamond Relaxometry-Based Detection of Free-Radical Species When Produced in Chemical Reactions in Biologically Relevant Conditions. ACS Sens 5, 3862–3869 (2020).

8. Sharmin, R. et al. Fluorescent Nanodiamonds for Detecting Free-Radical Generation in Real Time during Shear Stress in Human Umbilical Vein Endothelial Cells. ACS Sens 6, 4349–4359 (2021).

9. Wu, Y. & Weil, T. Recent Developments of Nanodiamond Quantum Sensors for Biological Applications. Advanced Science 9, 2200059 (2022).

10. Wu, K. et al. Applying NV center-based quantum sensing to study intracellular free radical response upon viral infections. Redox Biol 52, 102279 (2022).

11. Razinkovas, L., Maciaszek, M., Reinhard, F., Doherty, M. W. & Alkauskas, A. Photoionization of negatively charged NV centers in diamond: Theory and *ab initio* calculations. Phys Rev B 104, 235301 (2021).

12. Aslam, N., Waldherr, G., Neumann, P., Jelezko, F. & Wrachtrup, J. Photo-induced ionization dynamics of the nitrogen vacancy defect in diamond investigated by single-shot charge state detection. New J Phys 15, 013064 (2013).

13. Ninio, Y. et al. High-Sensitivity, High-Resolution Detection of Reactive Oxygen Species Concentration Using NV Centers. ACS Photonics 8, 1917–1921 (2021).

14. Gorrini, F. et al. Fast and Sensitive Detection of Paramagnetic Species Using Coupled Charge and Spin Dynamics in Strongly Fluorescent Nanodiamonds. ACS Appl Mater Interfaces 11, 24412–24422 (2019).

15. Pesah, Y. et al. Drosophila parkin mutants have decreased mass and cell size and increased sensitivity to oxygen radical stress. Development 131, 2183–2194 (2004).

16. Yang, Y. et al. Mitochondrial pathology and muscle and dopaminergic neuron degeneration caused by inactivation of Drosophila Pink1 is rescued by Parkin. Proc Natl Acad Sci U S A 103, 10793 (2006).

17. Shephard, F., Greville-Heygate, O., Marsh, O., Anderson, S. & Chakrabarti, L. A mitochondrial location for haemoglobins—Dynamic distribution in ageing and Parkinson’s disease. Mitochondrion 14, 64 (2014).

18. Stoll, S. & Schweiger, A. EasySpin, a comprehensive software package for spectral simulation and analysis in EPR. J Magn Reson 178, 42–55 (2006).

19. Sow, M. et al. Millikelvin Intracellular Nanothermometry with Nanodiamonds. Advanced Science e11670 (2025) doi:10.1002/ADVS.202511670.

20. Wu, Y. et al. Detection of Few Hydrogen Peroxide Molecules Using Self-Reporting Fluorescent Nanodiamond Quantum Sensors. J Am Chem Soc 144, 12642–12651 (2022).

21. Jarmola, A., Acosta, V. M., Jensen, K., Chemerisov, S. & Budker, D. Temperature- and Magnetic-Field-Dependent Longitudinal Spin Relaxation in Nitrogen-Vacancy Ensembles in Diamond. Phys Rev Lett 108, 197601 (2012).

22. Genov, G. T., Ben-Shalom, Y., Jelezko, F., Retzker, A. & Bar-Gill, N. Efficient and robust signal sensing by sequences of adiabatic chirped pulses. Phys Rev Res 2, 033216 (2020).

23. Allen, L., Eberly, J. H. & Physics. Optical Resonance and Two-Level Atoms (Dover Books on Physics). 256 (1987).

24. Wu, K. et al. Nanoscale detection and real-time monitoring of free radicals in a single living cell under the stimulation of targeting moieties using a nanodiamond quantum sensor. Functional Diamond 4, 2336524 (2024).

25. Cardoso Barbosa, I., Gutsche, J. & Widera, A. Impact of charge conversion on NV-center relaxometry. Phys Rev B 108, 075411 (2023).

26. Buettner, G. R. Spin trapping: ESR parameters of spin adducts. Free Radic Biol Med 3, 259–303 (1987).

27. Gan, Z. Y. et al. Activation mechanism of PINK1. Nature 602, 328–335 (2022).

28. Xiao, B. et al. Reactive oxygen species trigger Parkin/PINK1 pathway–dependent mitophagy by inducing mitochondrial recruitment of Parkin. J Biol Chem 292, 16697 (2017).

