## Supplementary Figures 1 to 3 for "Detecting Mitochondrial Free Radicals with Quantum Sensors: From Organelles to a *D. melanogaster* Model of Neurodegeneration"

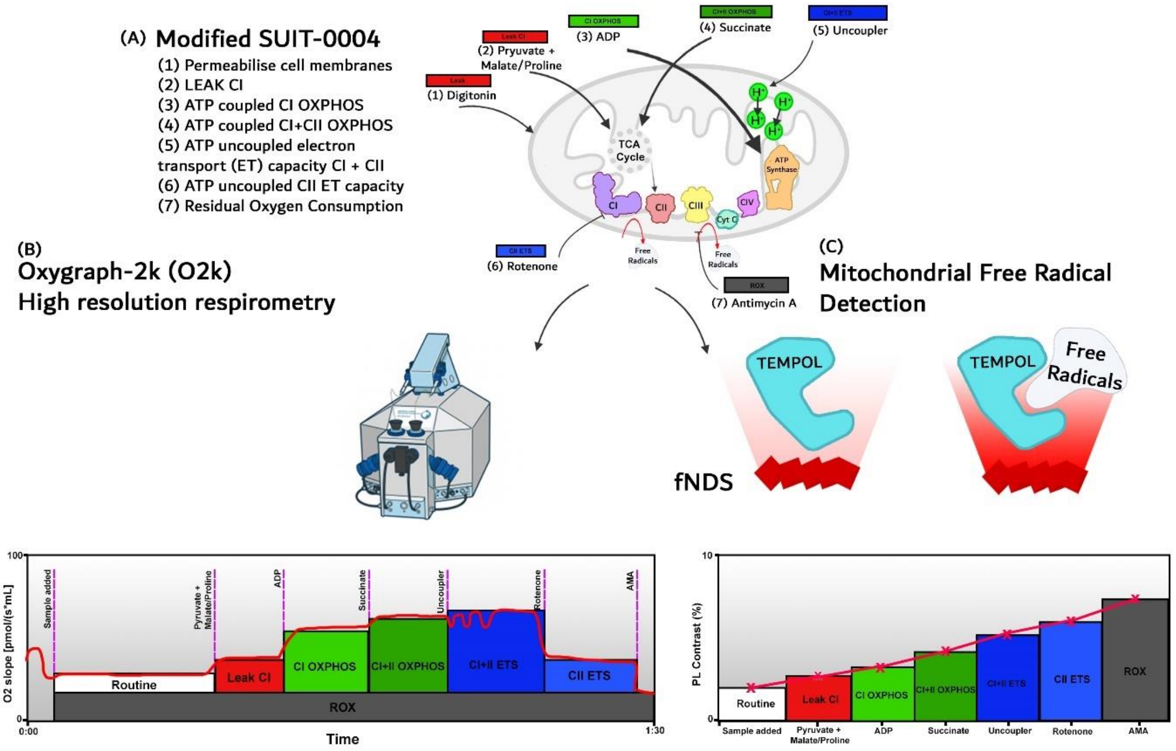


**Figure S1:** A visual representation of the key concepts of this technique. (A) The SUIT protocol sequentially probes mitochondrial complexes to assess changes in oxygen concentration using different substrates and inhibitors to target specific complexes. (B) Conventionally this protocol is used for the Oroboros Oxygraph-2k (O2k), a high-resolution respirometer that produces a trace akin to the schematic. The key points in which changes in oxygen consumption are anticipated are when the substrates are added, to which changes of free radical production due to electron leak from the ETS (Electron transport system) when undergoing oxidative phosphorylation (OXPHOS) are expected. (C) TEMPOL is used with the same protocol to be sensitive to these changes in mitochondrial respiration to scavenge free radicals when specific complexes are probed. TEMPOL sensitivity will reflect changes in free radical production at key points analogous to those described for isolated mitochondria, whole cells and whole organisms. An example trace is shown in which the addition of substrates and inhibitors gradually accumulates free radical formation leading to an increase in photoluminescence (PL) contrast.

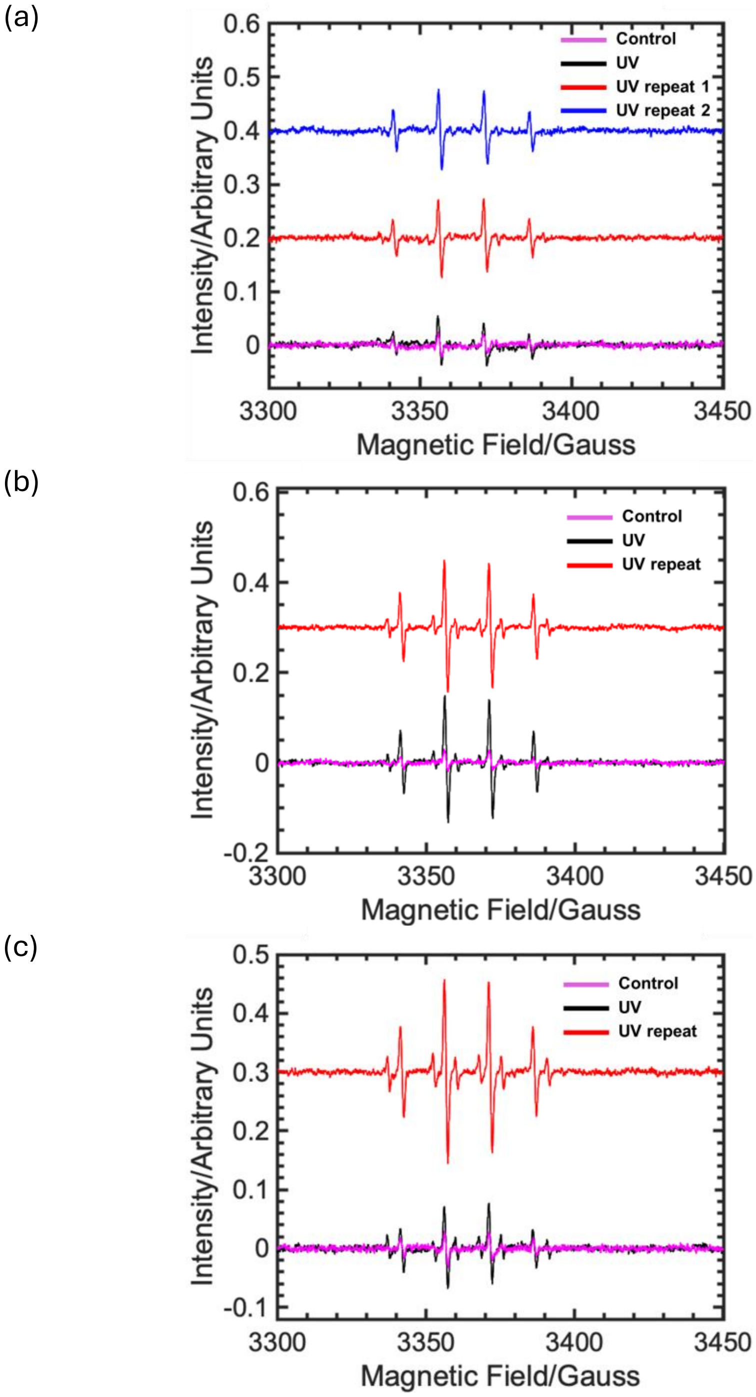


**Figure S2**: Continuous Wave (cw) EPR spectra of DMPO spin-trap adducts from $H_2O_2$ solutions at 30% (a), 20% (b), and 10% (c) concentrations. The magenta traces represent the un-irradiated controls, establishing a weak, low-amplitude baseline signal. In contrast, all traces recorded after UV-irradiation (black, red, blue) show increased signal amplitude, confirming UV-induced radical production. Critically, the overall signal intensity decreases with increasing $H_2O_2$ concentration, and the relative proportion of 'sextet' to 'quartet' species also decreases with increasing $H_2O_2$ concentration. The red traces (all panels) and the blue trace (30% panel only) are repeat measurements taken ~1 hour after the initial scan (black trace). While the repeats in the 30% and 20% panels show similar amplitudes, demonstrating good reproducibility, the notably different amplitude between the black and red traces in the 10% panel indicates significant sample instability over this period. Traces are vertically offset for clarity.

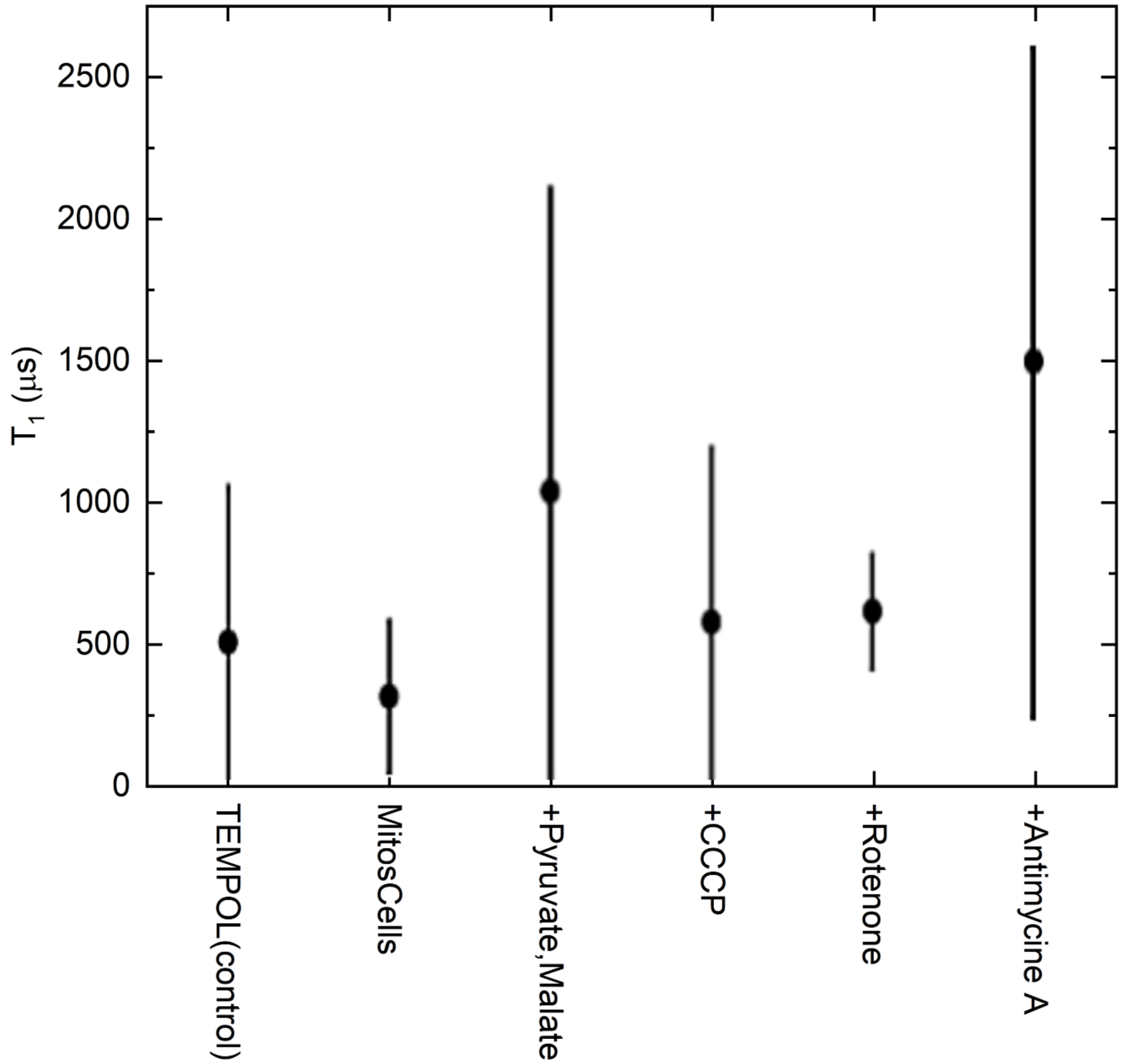


**Figure S3**: Nanodiamond $T_1$ relaxation times at various stages of the Substrate-Uncoupler-Inhibitor Titration (SUIT) protocol. Error bars represent the standard deviation obtained from fitting the $T_1$ measurement data to a stretched exponential function.
